# A cell-permeant nanobody-based degrader that induces fetal hemoglobin

**DOI:** 10.1101/2022.06.07.495197

**Authors:** Fangfang Shen, Ge Zheng, Mekedlawit Setegne, Karin Tenglin, Manizheh Izada, Henry Xie, Liting Zhai, Stuart H. Orkin, Laura M. K. Dassama

## Abstract

Proximity-based strategies to degrade proteins have enormous therapeutic potential in medicine, but the technologies are limited to proteins for which small molecule ligands exist. The identification of such ligands for therapeutically relevant but “undruggable” proteins remains challenging. Herein, we employed yeast surface display of synthetic nanobodies to identify a protein ligand selective for BCL11A, a critical repressor of fetal globin gene transcription. Fusion of the nanobody to a cell-permeant miniature protein and an E3 adaptor creates a degrader that depletes cellular BCL11A in erythroid precursor cells, thereby inducing the expression of fetal hemoglobin, a modifier of clinical severity of sickle cell disease and β-thalassemia. This work establishes a new paradigm for the targeted degradation of previously intractable proteins using cell-permeant nanobody-based degraders.

**One sentence summary:** A cell-permeant, protein-based degrader is used for the induction of fetal hemoglobin.

## Main text

Proteolysis targeting chimeric molecules (PROTACs) and molecular glue degraders, such as the immunomodulatory imide drugs, hijack the cellular protein ubiquitination machinery to specifically degrade proteins of interest (POIs) (*1, 2*). They offer exciting opportunities for use as therapeutics and powerful research tools for biological inquiry (*3, 4*). While PROTAC molecules are being used clinically with notable success and great promise (*5*), it remains challenging to target many therapeutically relevant protein families due to challenges inherent in ligand discovery and optimization. This is especially true for transcription factors (TFs), which generally contain unstructured domains and lack obvious “ligandable” pockets (*6, 7*). Additionally, the development of PROTACs requires substantial synthetic efforts to test various combinations of recruited ubiquitin E3 ligases and linkers that are optimal for forming a ternary complex of the PROTAC, POI, and ubiquitin E3 ligase (*8*). Other elegant degradation platforms, including the degradation tag system (*9*) and transcription factor targeting chimeras (TRAFTACs) (*10*), have been developed for difficult protein targets. However, the utility of these platforms is either limited to engineered proteins or their specificity for the targeted proteins unexplored.

Unlike the small molecule ligands that are typically used in PROTACs, antibodies exploit features of protein surfaces to recognize antigens, and show exceptional specificity and remarkable affinity for their antigens. Even fragments of single variable heavy chain domains, termed nanobodies (Nbs), retain antigen specificity and can be used as the POI ligand in a PROTAC. The recent development of a yeast surface display platform to screen large libraries of synthetic nanobodies provides a straightforward and low-cost method to obtain Nb ligands for proteins (*11*). While protein-based degraders using Nbs (*12-17*) or the Fc domain of antibodies (Trim-Away) (*18*) have been reported, their potential is hindered either by the challenge of delivering these ligands to intracellular targets or because the ligands do not target endogenous proteins. We hypothesized that appending a cell penetrating moiety to Nb degraders could overcome these limitations and provide a broad strategy to target endogenous proteins. To demonstrate the utility of this approach, we focused on the transcription factor BCL11A, because it is a clinically-validated target for the treatment of hemoglobin disorders, including β-thalassemia and sickle cell disease (SCD) (*19, 20*).

Reactivation of fetal hemoglobin (HbF, α_2_γ_2_) is a promising strategy to ameliorate clinical severity in patients with hemoglobin disorders (*21*). Patients with SCD that produce elevated HbF exhibit significantly improved survival rates (*22*). BCL11A represses HbF expression through direct binding at the γ-globin promoters, eliciting the switch from fetal to adult hemoglobin (HbA, α_2_β_2_) expression during erythropoiesis (*19, 23, 24*). Genetic approaches, notably clustered regularly interspaced short palindromic repeats (CRISPR)–Cas9 (*25, 26*) editing and RNA interference (*27*) have been used to downregulate BCL11A in patients and validate the clinical utility of disabling BCL11A. However, the resource-intensive care and high cost of ex vivo genetic manipulation of cells in these trials limit the application of these treatments. PROTACs may provide an alternative means to deplete BCL11A in a temporal, reversible, precise, and cost-effective manner.

Although BCL11A is predicted to be largely unstructured, the protein contains several well-ordered regions, including a CCHC-type zinc finger domain (ZnF0) that might mediate self-association (*28*) and six C_2_H_2_-type zinc finger domains (ZnF1, ZnF23 and ZnF456) (Fig. 1a). We reasoned that such well-folded domains could be used to identify Nb ligands for BCL11A. Because of the close sequence similarity between BCL11A and its paralog BCL11B in all of the zinc finger regions (*29*), ligands that bind to these domains might demonstrate affinity for both paralogs. With the aim of achieving specificity in ligands selected for BCL11A, we produced ZnF23 and an “extended” zinc finger domain (exZnF23), which includes the C-terminal unstructured region that diverges in sequence between the two paralogs (Fig. 1a). The protein fragment was expressed in *E. Coli* and purified. We used recombinant protein in a screen using a yeast surface display platform to identify synthetic Nbs that bind to exZnF23 (Fig. 1b). An initial Nb hit (**wt 2D9**) was produced in *E. coli* (Fig. S1a) and its affinity for BCL11A was assessed using a pull-down assay (Fig. S1b). MicroScale Thermophoresis (MST) measurements further revealed that **wt 2D9** binds specifically to BCL11A exZnF23, and not to ZnF23 or BCL11B exZnF23, despite the high similarity in the Zn finger domains (Fig. S1c). Affinity maturation through random mutagenesis of **wt 2D9** was performed, after which 7 Nbs with better affinities were obtained (Fig. S2). Of these, **2D9_V102G** (hereafter referred to as **2D9**) and **2D9_W108L** were selected for additional studies because of their stability, high affinity, and specificity for exZnF23 of BCL11A (Figs. 1c, S3). Size-exclusion chromatography coupled with multi-angle light scattering revealed that BCL11A exZnF23 is monomeric and formed a stable complex with **2D9** (Fig. S4). A crystal of **2D9** complexed with exZnF23 of BCL11A was obtained, but insufficient electron density precluded modeling of BCL11A. Nonetheless, the high-resolution structure of **2D9** combined with maturation mutagenesis data suggested that some of the interaction with BCL11A is mediated through a loop in Complementarity Determining Region (CDR) 3 of the Nb (Fig. S5).

**Figure 1.**
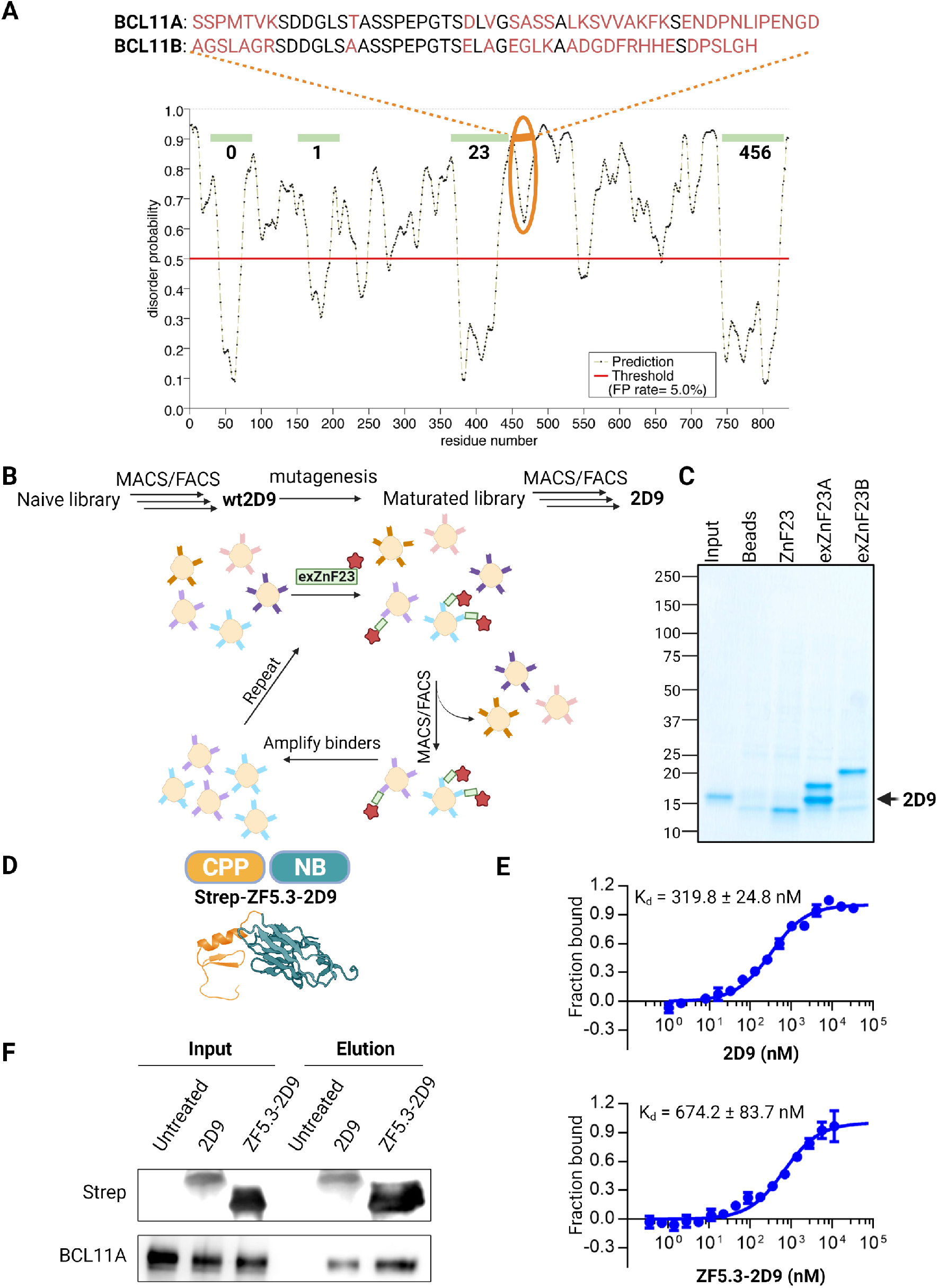
Identification and construction of cell permeant Nb ligands for BCL11A. (**A**) Disorder probability of BCL11A revealing predicted order in the ZnF domains (**0-6**) and the sequence divergence from BCL11B in the extended ZnF23 region. The ZnF23 domains of the paralogs are 93.2% identical and 96.6% similar; in the exZnF23 fragment, the sequence identity and similarity are 69.3% and 78.1%, respectively. (**B**) Flow chart summarizing Nb selection. (**C**) SDS-PAGE analysis showing that Nb **2D9** interacts with exZnF23 of BCL11A (exZnF23A) but not ZnF23 and exZNF23 of BCL11B (exZnF23B). (**D**) Schematic description of the cell-permeant Nb **ZF5.3-2D9** created from the individual structures of ZF5.3 (modeled secondary structure), and **2D9** (PDB 7UTG). (**E**) MST analysis of the binding of **2D9** and **ZF5.3-2D9** to exZnF23 of BCL11A. (**F**) Immunoprecipitation revealing that **2D9** and **ZF5.3-2D9** bind to endogenous BCL11A.

Ligands intended for depletion of BCL11A protein must be delivered to the nucleus of erythroid precursor cells. Because Nbs are too large and unfavorably charged to traverse the plasma membrane, **2D9** was functionalized for cell penetration by appending a cell permeant miniature protein called ZF5.3 (*30-32*) (Fig. 1d). MST measurements demonstrated that appending ZF5.3 to **2D9** did not significantly alter the affinity of **2D9** for BCL11A exZnF23 (Fig. 1e). Addition of purified **2D9** or the fusion protein **ZF5.3-2D9** to the lysate of human umbilical cord blood-derived erythroid progenitor (HUDEP-2) cells (*33*) confirmed their association with endogenous, full-length BCL11A (Fig. 1f). Additional experiments in which **ZF5.3-2D9** was incubated with HUDEP-2 cells resulted in accumulation of a significant fraction of the protein localized to the nucleus, as evidenced by confocal imaging of immunofluorescent stained cells (Fig. 2a). Immunoblotting demonstrated a dose-(Fig. 2b) and time-dependent cellular uptake (Fig. 2c), and cell fractionation revealed that the fusion protein was present in the nucleus for up to 24 hrs (Fig. 2d). Co-immunoprecipitation of BCL11A from HUDEP-2 cells incubated with **ZF5.3-2D9** further revealed its cellular entrance and binding to BCL11A (Fig. 2e).

**Figure 2.**
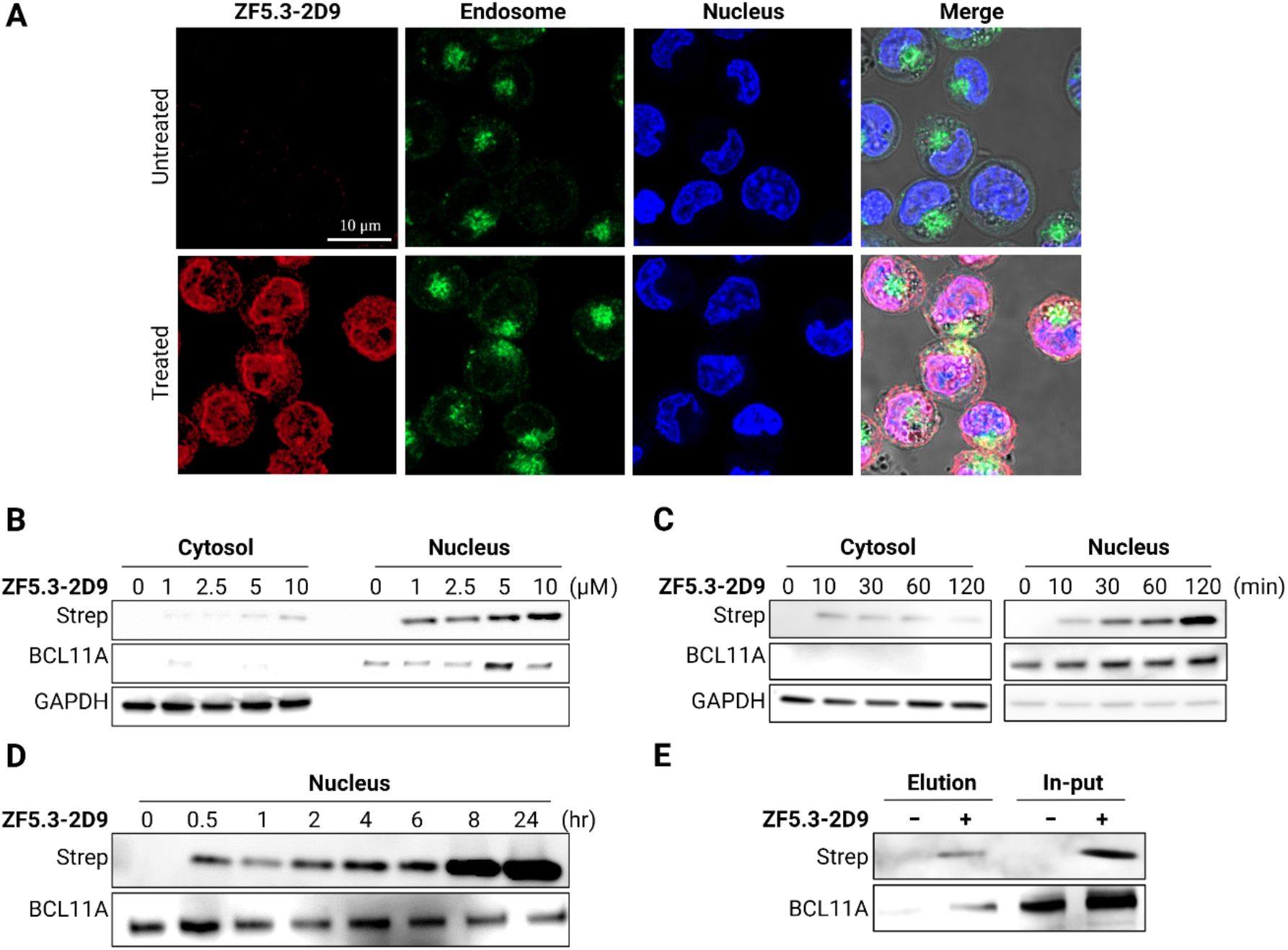
Delivery of **ZF5.3-2D9** to erythroid precursor cells. (**A**) Confocal microscopy images of HUDEP-2 cells revealing **ZF5.3-2D9** (red) entered HUDEP-2 cells and show significant colocalization (purple) with the nucleus (blue). Immunoblots revealing a (**B**) concentration- and (**C**) time-dependent cell penetration by **ZF5.3-2D9**. (**D**) Immunoblot showing increased accumulation of **ZF5.3-2D9** in the nucleus. (**E**) Co-immunoprecipitation of endogenous BCL11A using delivered **ZF5.3-2D9**. GAPDH and BCL11A were used as loading controls for the cytosolic and nuclear fractions, respectively.

Having demonstrated that **ZF5.3-2D9** penetrated HUDEP-2 cells and engaged BCL11A, we used the cell-permeant Nb as a handle to mediate the proteasomal degradation of BCL11A. The rational design of PROTACs is difficult due to an incomplete understanding of rules that govern formation of the ternary complex between the POI, ubiquitin E3 ligase, and the PROTAC. Unlike small molecule PROTACs, “all protein” degraders using reengineered E3 ligases are reported to exhibit high flexibility to various targets (*12, 15*). However, these ligands are rarely cell permeant and do not degrade endogenous proteins, thereby limiting their utility.

To first confirm that **2D9** can mediate selective degradation of BCL11A, we produced plasmids of the Nb fused to the Fc domain of Immunoglobulin G1 (Nb-Fc) or Trim 21 (Nb-wtTrim21). With these designs, TRIM 21 would mediate proteasomal degradation via the Trim-Away method (*18*). Lentiviral transduction of HUDEP-2 cells with these plasmids revealed that all constructs induced loss of BCL11A and reactivation of HBG (Fig. S6). Similar experiments performed with a Trim 21 variant (2D9-mutTrim21 or W108L-mutTrim21) revealed no change in BCL11A levels. To assess the *in vivo* specificity of the Nbs for BCL11A, we overexpressed BCL11B in HEK293T cells and transfected the cells with Nb-Fc and Nb-wtTrim21; neither of promoted loss of BCL11B (Fig. S7). Together, these data indicate that the Nbs **2D9** and **2D9_W108L** can distinguish BCL11A from BCL11B and target the endogenous BCL11A with high specificity.

Encouraged by these results, we designed cell-permeant, nanobody-based degraders for BCL11A by incorporating two ubiquitin E3 ligases into our design: SPOP (speckle type POZ protein) and RNF4. These E3 ligases were chosen because of their high abundance in the nucleus and their previous use to mediate the degradation of nuclear proteins (*13, 17*). SPOP is an E3 adaptor protein that functions in complex with cullin-3 (CUL3); it is composed of a substrate binding MATH domain and a CUL3-binding BTB domain. RNF4 is an E3 ligase that contains an N-terminal SUMO substrate binding site and a C-terminal RING domain responsible for dimerization and E2 binding. By replacing the native substrate recognition domain of SPOP and RNF4 with **ZF5.3-2D9**, we created the proteins **ZF5.3-2D9-SPOP** and **ZF5.3-2D9-RNF4** (Fig. 3a). Both proteins were expressed and purified from *E. coli* (Fig. S8), and were delivered in pure form to HUDEP-2 cells through incubation (Fig. S9). Upon their delivery, BCL11A levels were lowered steadily over time (Fig. 3b, S10). With **ZF5.3-2D9-SPOP**, the loss was striking, as up to 70% of BCL11A was depleted within 12 hours of incubation with 10 μM of the degrader (Fig. 3c). Because of its far greater degradation effect than **ZF5.3-2D9-RNF4, ZF5.3-2D9-SPOP** was used for subsequent studies. With this degrader protein, loss of BCL11A in HUDEP-2 cells was sustained for at least 72 hours (Fig. S11). In control experiments with **ZF5.3-SPOP**, we verified that BCL11A loss required **2D9** (Fig 3d). Treatment with a proteasome inhibitor (**MG-132**) prevented the degradation of BCL11A, thereby confirming that the loss of the protein proceeded via proteasomal degradation (Fig. 3e).

**Figure 3.**
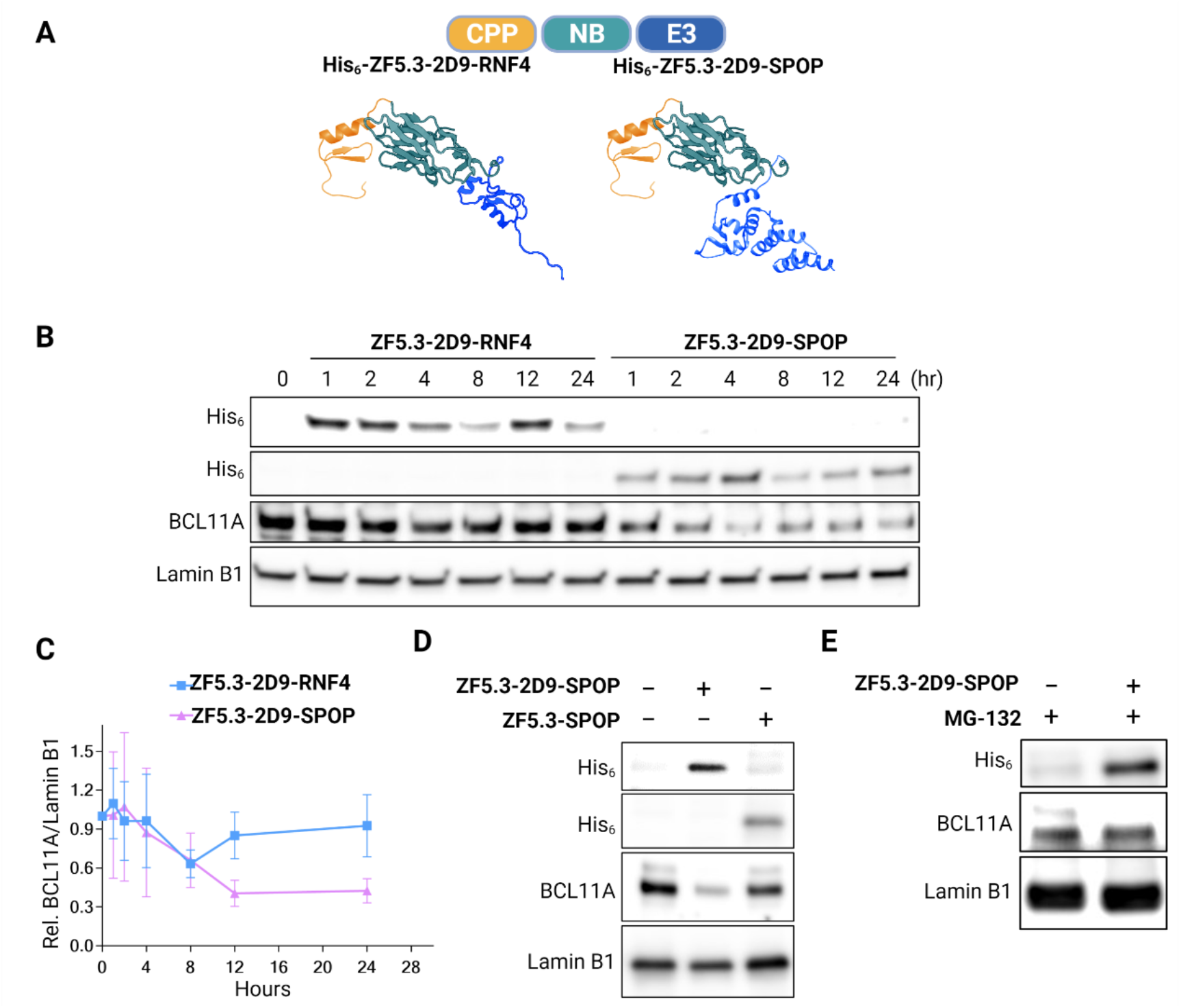
Nanobody-mediated degradation of BCL11A in HUDEP-2 cells. (**A**) Schematic depiction of BCL11A degraders. The individual structures of the RNF4 RING domain (4PPE), the BTB domain of SPOP (3HTM), ZF5.3 (modeled secondary structure), and **2D9** (7UTG) were retrieved from the Protein Data Bank. (**B**) Representative immunoblot showing loss of BCL11A in HUDEP-2 cells treated with **ZF5.3-2D9-RNF4** or **ZF5.3-2D9-SPOP**. (**C**) Quantification of BCL11A loss from three independent biological replicates. Immunoblots revealing that (**D**) loss of BCL11A requires the presence of **2D9** and (**E**) is prevented by addition of 5 μM of the proteasome inhibitor **MG**-**132**. Lamin B1 was used as a loading control for the immunoblots.

Endogenous expression of BCL11A in HUDEP-2 cells varies during differentiation (Fig. S10) and represses expression of the fetal-stage γ-globin genes. Given the sustained loss of BCL11A in response to treatment with the degrader in undifferentiated HUDEP-2 cells, we explored its effect on differentiated HUDEP-2 cells. HUDEP-2 cells were treated with **ZF5.3-2D9-SPOP** on Days 0 and 3 of differentiation, and samples were collected on Days 3 (prior the second treatment), 4 and 7 for analysis (Fig. 4a). RT-qPCR of hemoglobin transcripts in samples from Day 7 (Fig. 4b) revealed an increase of γ-globin transcripts in HUDEP-2 cells treated with **ZF5.3-2D9-SPOP**. Immunoblots revealed that the degradation of BCL11A was maintained throughout and that significant fetal hemoglobin (HbF) was induced by Day 4 (Fig. 4c). HbF immunostaining and fluorescence activated cell sorting of HUDEP-2 cells provided additional confirmation that treatment with **ZF5.3-2D9-SPOP** resulted in a 2.5-fold increase of the HbF^+^ population (Fig. 4d).

**Figure 4.**
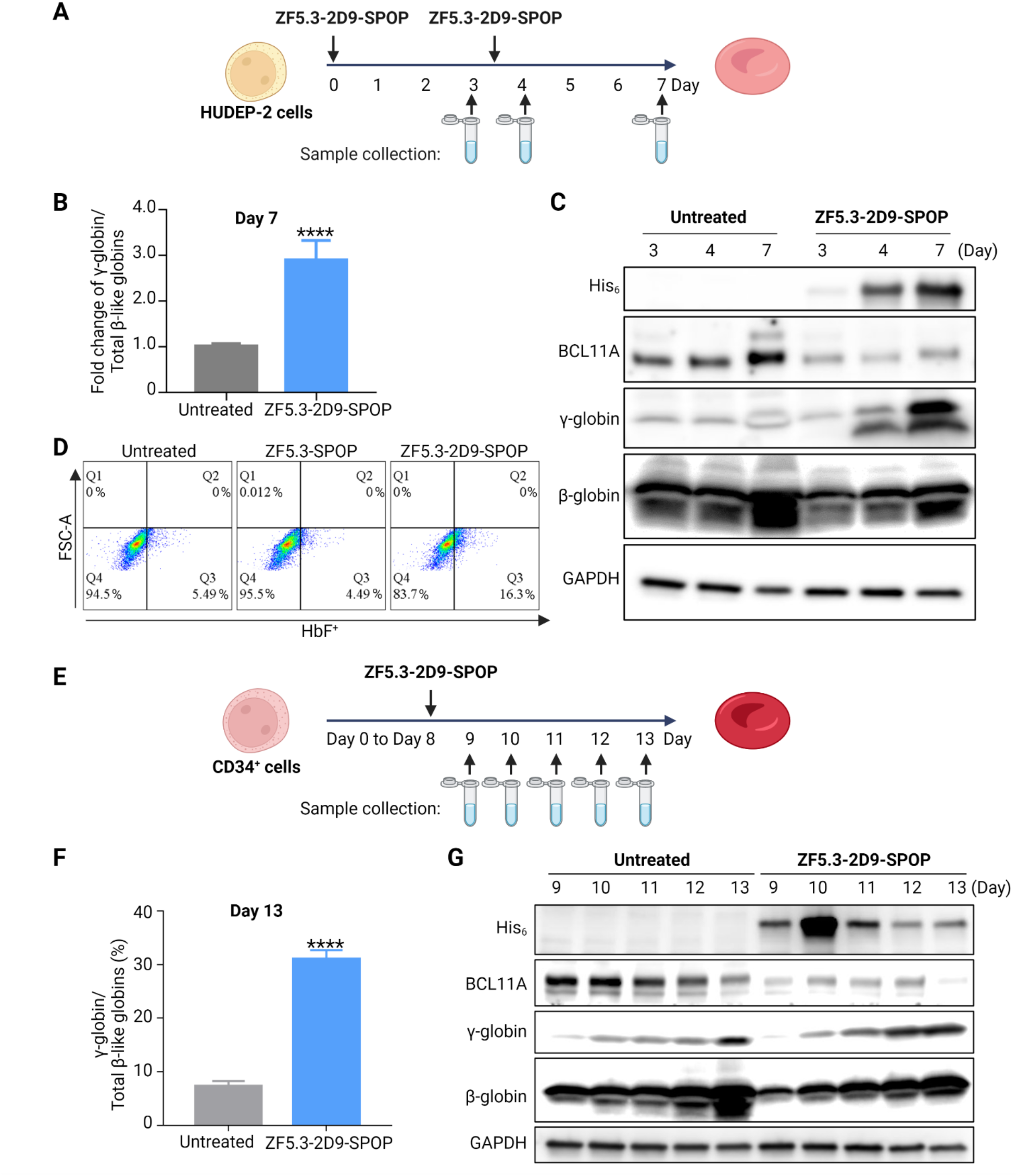
HbF induction in HUDEP-2 and CD34^+^ cells. (**A**) Schematic depiction of the HUDEP-2 cell differentiation and treatment. (**B**) qRT-PCR showing an increase in γ-globin mRNA after treatment with **ZF5.3-2D9-SPOP** in HUDEP-2 cells. (**C**) Immunoblot revealing the loss of BCL11A and γ-globin increase after treatment with **ZF5.3-2D9-SPOP** in HUDEP-2 cells. (**D**) FACS analysis of immunostained HUDEP-2 cells from Day 7 of differentiation showing an increase in the population of HbF-positive cells following the degradation of BCL11A. (**E**) Schematic depiction of the CD34^+^ cell differentiation and treatment. (**F**) qRT-PCR showing increase of γ-globin mRNA level after treatment with **ZF5.3-2D9-SPOP** in CD34^+^ cells. (**G**) Immunoblot revealing the loss of BCL11A and γ-globin increase after treatment with **ZF5.3-2D9-SPOP** in CD34^+^ cells. GAPDH was used as a loading control in all immunoblots.

We further investigated the effect of **ZF5.3-2D9-SPOP** in human primary CD34^+^ progenitor cells. The cells were cultured under differentiation conditions (Fig. S12) and treated with **ZF5.3-2D9-SPOP** on Day 8 of the differentiation (Fig. 4e). As with HUDEP-2 cells, immunoblots of samples from Days 9-13 revealed sustained degradation of BCL11A and marked HbF induction as of Day 11 (Fig. 4f). RT-qPCR of hemoglobin transcripts in samples from Day 13 showed an increase of γ-globin to 30% of the total beta-like hemoglobin, a level above the threshold for inhibition of red blood cells sickling SCD patients (*22*) (Fig. 4g). Together, these data provide evidence that the degrader **ZF5.3-2D9-SPOP** depletes BCL11A and elicits increased expression of HbF in erythroid precursor cells.

In conclusion, this work presents a systematic strategy for modulating the function of a traditionally “undruggable” target. We employed strategies to identify a selective nanobody ligand of BCL11A by directing its binding to an unstructured region of the protein that shares low similarity with the closest paralog, BCL11B. The Nb ligand was functionalized for cell penetration and used to construct a degrader for the proteasomal degradation of BCL11A. If methods for efficient delivery of Nbs to cells *in vivo* can be developed, the degrader strategy described here could be considered for treatment of SCD and β-thalassemia. Apart from any clinical application, these studies establish a paradigm through which disease-relevant but intractable proteins can be targeted for modulation for function validation or biological studies. The use of a readily available nanobody library, genetically encoded cell-penetrating vehicle, and an engineered E3 adapter protein significantly simplifies efforts to degrade other “undruggable” proteins.

## Acknowledgements

We thank Andrew Kruse for providing the yeast display library and Ms. Lisha Ou and Mr. Jonathan Chou for helpful discussions.

## Author contributions

F.S., G.Z., S.H.O., and L.M.K.D. designed the study; F.S., G.Z., M.S., K.T., M. I., H. X., and L. Z. performed research, analyzed data, and prepared figures; F.S., G.Z., S.H.O., and L.M.K.D. wrote the manuscript.

## Data availability

The crystallographic structure for 2D9 has been deposited in the Protein Data Bank (PDB 7UTG). All other data are available in the manuscript or supplementary materials.

## Supplementary Materials

Materials and Methods

Figures S1 – S13

Tables S1 – S6

