## Supplementary materials for "A cell-permeant nanobody-based degrader that induces fetal hemoglobin"

**This PDF file includes:**

Materials and Methods

Supplemental Figures S1 – S13

Supplemental Tables S1– S6

### Materials and Methods

***Cell culture.*** HUDEP-2 cells (RCB4557) were obtained from Riken BioResource Research Center (Japan). Cells were cultured according to the reported method *(33)*. In brief, cells were maintained in expansion medium, which contains StemSpan serum-free expansion medium (SFEM, Stemcell Technologies), 2% Penicillin-Streptomycin solution (10,000 U/mL stock), 50 ng/mL recombinant human stem cell factor (SCF, Stemcell Technologies), 3 IU/mL Epoetin alfa (Epogen, Amgen), 0.4 μg/mL dexamethasone, and 1 μg/mL doxycycline. For differentiation, cells were transferred from expansion medium to EDM-2, which contains Iscove's modified Dulbecco's medium (IMDM), 1% L-glutamine (this is in addition to the L-glutamine present in IMDM), 2% Penicillin-Streptomycin solution (10,000 U/mL stock concentration), 330 μg/mL human holo-transferrin, 10 μg/mL recombinant human insulin solution, 2 IU/mL heparin, 5% inactivated human plasma type AB, 3 IU/mL Epoetin alfa, 100 ng/mL SCF and 1 μg/mL doxycycline. After culturing for 4 days, cells were transferred to EDM-3 (same as EDM-2 but without SCF) and cultured for another 3 days. After that, cells were cultured in EDM (no doxycycline) for 2 days.

Human HEK293T cells (female) were purchased from ATCC. The cells were cultured in DMEM, high glucose (Thermo Fisher Scientific, 11965) with 10% FCS and 2 mM L-Glutamine.

Peripheral blood derived $CD34^+$ cells were obtained from the NIDDK-Center of Excellence in Hematology at the Fred Hutchinson Cancer Research Center. The cells were cultured according to the reported method (*34*). In brief, cells were cultured in erythroid differentiation medium (EDM) which contains IMDM supplemented with stabilized glutamine, 330 ug/mL holo-human transferrin, 10 ug/mL recombinant human insulin, 2 IU/mL heparin Choay, and 5% inactivated human plasma type AB. The expansion procedure comprised 3 steps. In the first step (Day 0 to Day 7), $10^4$/mL $CD34^+$ cells were cultured in EDM in the presence of $10^{-6}$ M hydrocortisone (Stemcell Technologies), 100 ng/mL SCF, 5 ng/mL IL-3 (Stemcell Technologies), and 3 IU/mL Epoetin alfa. On Day 4, 1 volume of cell culture was diluted in 4 volumes of fresh medium containing SCF, IL-3, Epoetin alfa, and hydrocortisone. In the second step (Day 7 to Day 11), the cells were resuspended at $10^5$/mL in EDM supplemented with SCF and Epo. In the third step (Day 11 to Day 18), the cells were cultured in EDM supplemented with Epo alone. Cell counts were adjusted to a range of $7.5 \times 10^5$ - $1 \times 10^6$ on Day 11. Beyond Day 18, the culture medium containing Epo was renewed twice a week.

***Plasmid construction***. All gene blocks were obtained from GENEWIZ while primers were obtained from Integrated DNA Technologies. KOD hot start polymerase was used for PCR reaction, NEBuilder HiFi DNA Assembly was used for plasmid fusion. NdeI, Xhol and T4 ligase were purchased from New England Biolabs.

<u>pET-20b_2D9 Plasmid Construction</u>

The sequence of 2D9 with a N-terminal Strep-tactin tag was codon-optimized for expression in *E. coli* and cloned into a linearized pET-20b vector at the NdeI and XhoI restriction sites.

<u>pET-20b_ZF5.3-2D9 Plasmid Construction</u>

The sequence of ZF5.3 was codon optimized for expression in *E. coli* and cloned into a linearized pET20b_2D9 plasmid with a N-terminal Strep-tactin tag.

<u>pET-28a ZF5.3-2D9 -SPOP Plasmid Construction</u>:

The sequence of $SPOP_{167-374}$ was codon-optimized for expression in *E. coli* and cloned into a linearized pET-20b_2D9 plasmid with a N-terminal Strep-tactin tag. After the HiFi assembly, pET-20b_ZF5.3-2D9-SPOP was digested with restriction enzymes NdeI and XhoI and ligated into pET-28a containing a N-terminal $His_6$ tag to obtain pET-28a_ZF5.3-2D9-SPOP plasmid.

pET-20b_ZF5.3-2D9-RNF4 Plasmid Construction:

The sequence of $RNF4_{75-194}$ was codon-optimized for expression in *E. coli* and cloned into a linearized pET-20b_2D9 plasmid with a N-terminal Strep-tactin tag. Primers used are summarized in Table S1.

***Expression and purification of proteins ZnF23, exZnF23 of BCL11A and exZnF23 of BCL11B***. The cDNA of ZnF23 (residues 372-430) and exZnF23 (residues 372-484) of human BCL11A, exZnF23 of human BCL11B (residues 422-528) were cloned into pET28a vector and expressed as N-terminal $His_6$-tag fusion proteins in *E. coli*. Cells were resuspended in lysis buffer (50 mM Tris-HCl pH 8.0, 300 mM NaCl, 10 mM imidazole, 1 mM DTT). After sonication and centrifugation, the supernatant was applied to the $Ni^{2+}$-NTA resin (Qiagen) equilibrated with lysis buffer and incubated at 4°C for 1 hr. The resin was washed with wash buffer I (50 mM Tris-HCl pH 8.0, 1 M NaCl, 10 mM imidazole, 1 mM DTT), then wash buffer II (50 mM Tris-HCl pH 8.0, 300 mM NaCl, 10 mM imidazole, 1 mM DTT), and stepwise eluted with elution buffer (50 mM Tris-HCl pH 8.0, 300 mM NaCl, 100/200/300/500 mM imidazole, 1 mM DTT). The eluate was examined by SDS-PAGE stained with Coomassie blue. Fractions containing the target proteins were combined and then dialyzed with dialysis buffer (1×PBS, 150mM NaCl, 2mM DTT) at 4 ℃ overnight and concentrated before loading onto the HiLoad 16/600 Superdex 75 prep-grade column (Cytiva) equilibrated with the purification buffer (1×PBS, 1 mM DTT). Purified proteins were concentrated, flash frozen in liquid nitrogen and stored at -80°C for yeast screening and binding assays.

***Isolation of BCL11A-exZnF23-binding nanobodies from yeast synthetic library***. Purified exZnF23 of BCL11A protein was labeled with AlexaFluor647 dye (Invitrogen) or fluorescein isothiocyanate (FITC) (Sigma-Aldrich) according to the manufacturer's protocols.

For the first round of magnetic-activated cell sorting (MACS), $1x10^{10}$ *S. cerevisiae* cells expressing a surface displayed library of synthetic nanobodies *(11)* were centrifuged, resuspended in selection buffer (20 mM HEPES pH 7.5, 150 mM NaCl, 0.1% BSA, 5 mM maltose) and then incubated with anti-AlexaFluor647 microbeads (Miltenyi) at 4°C for 40 min. The yeast cells were then passed through an LD column (Miltenyi) to remove any yeast expressing nanobodies that non-specifically interacted with the microbeads. Yeast cells that flowed through the column were centrifuged, resuspended in selection buffer, and incubated with 1 μM of AlexaFluor647-labeled exZnF23 of BCL11A at 4°C for 1 hr. Yeast cells were then centrifuged, resuspended in selection buffer with anti-AlexaFluor647 microbeads, and incubated at 4°C for 20 min before passing through an LS column (Miltenyi). The eluted yeast cells were collected and expanded to a subsequent round of MACS to further enrich for exZnF23-binding nanobodies. The second round of MACS was performed similarly to the first round but using fluorescein isothiocyanate (FITC)-labeled exZnF23 of BCL11A and anti-FITC microbeads. After MACS selections, yeast cells were co-stained with AlexaFluor647- and FITC-labeled exZnF23 of BCL11A and sorted by flow cytometry (Sony SH800Z). Double-positive yeast cells were selected and plated as single colonies, which were randomly picked and grown as clonal populations in 96-well plates. Yeast cells in 96-well plates were induced, stained with AlexaFluor647- or FITC-labeled exZnF23 of BCL11A and analyzed by the plate reader. Yeast DNA was extracted using standard methods and sequenced from the high activity clones.

***Affinity maturation of nanobody wt2D9***. Error-prone PCR was performed on nanobody wt2D9 DNA using the GeneMorph II Random Mutagenesis Kit (Agilent) and the resulting library was scaled up with a second PCR using Q5 High-Fidelity DNA Polymerase (New England Biolabs). 100 mL of BJ5465 *S.cerevisiae* cells were grown to $OD_{600nm}$ of 1.8 and were then made into electrocompetent yeast cells with 100 mM lithium acetate (*35*). The electrocompetent cells were transformed with 56 μg of the error prone library and 17 μg of linearized pYDS649 plasmid *(11)* using an ECM 830 Electroporator (BTX-Harvard Apparatus) with 500 V and 15 ms single pulse. The resulting library of nanobody 2D9 mutants has a mean mutation rate of about 1 amino acid change per nanobody clone.

$1\times10^6$ yeast cells from the error prone library were stained with anti-HA AlexaFluor488 antibody in selection buffer (20 mM HEPES pH 7.5, 150 mM NaCl, 0.1% BSA, 5mM maltose, 1mM DTT) to assess nanobody expression levels. In order to obtain high-affinity binders to exZnF23 of BCL11A, 4 rounds of MACS selections were performed, and the yeast cells were stained with 1μM of FITC-labeled exZnF23, 1μM of FITC-labeled exZnF23, 1.5μM of $His_6$-SBP-exZnF23, and 1 μM of FITC-labeled exZnF23 to enrich for binders with higher affinities. After MACS selections, 2 rounds of FACS were performed. In the first round of FACS, the yeast cells were co-stained with 0.5 μM of AlexaFluro647-labeled exZnF23 and 0.75μM of FITC-labeled exZnF23. A total of ~80,000 yeast cells from the first round of FACS were expanded and used for a second round to further enrich for high affinity nanobodies. In second round of FACS, the yeast cells were co-stained with 0.1 μM of AlexaFluro647-labeled exZnF23 and 0.15 μM of FITC-labeled exZnF23. After the second round of FACS, approximately 5,000 yeast cells were plated as single colonies using serial dilutions. 96 yeast clones were randomly selected, mini-prepped and sequenced to reveal consensus mutations contributing to affinity. The sequence of wt2D9 is found in Table S2.

***Nanobody degrader production***

Transformation of plasmids in BL21 cells.

2 μL of plasmid (concentration ≥20 ng/μL) containing either 2D9, ZF5.3-2D9, ZF5.3-2D9 -SPOP, ZF5.3-2D9 -RNF4, or ZF5.3-SPOP was added to a thawed tube of *E. coli* BL21 cells and incubated on ice for 15 min. The cells were heat shocked at 42 °C for 45 s and placed on ice for 2 min. 900 μL of LB media were added and the cells cultured for 1 h at 37 °C while shaking at 200 rpm. Cells were centrifuged for 3 min at 2500 × g and 900 μL LB media was removed. The recovered cells were resuspended in 100 μL remaining media and added to ampicillin-(100 μg/mL) or kanamycin-containing (50 μg/mL) agar plates and incubated at 37 °C overnight.

2D9 expression and purification

The plasmid encoding Strep-2D9 was used to transform *E. coli* BL21 cells. Individual colonies were selected on the basis of Amp resistance and used to inoculate 50 mL of LB media supplemented with Amp (100 mg/L). The primary culture was grown overnight and then used to inoculate 6 L of ZYM-5052 autoinduction (AI) media (*36*) supplemented with ampicillin, which was then allowed to grow at 37 °C with shaking at 200 rpm. When the $OD_{600nm}$ reached 0.5, the temperature was changed to 18 °C and the cells cultured overnight. The cells were harvested by centrifugation at 6000 × g for 30 min at 4 °C, resuspended in buffer containing 25 mM Tris-HCl pH 8.0, 300 mM NaCl, and 10% glycerol, supplemented with 1 mM PMSF and lysed via microfluidization. The lysate was clarified by centrifuging at 15,000 × g for 30 min at 4 °C and the cleared lysate manually added to a column with 5 mL Strep-Tactin® Sepharose® resin (IBA Lifesciences). The column was washed with 5 column volumes of 25 mM Tris-HCl pH 8.0, 300 mM NaCl, and 10% glycerol. Then protein was eluted from the resin using 25 mM Tris-HCl pH 8.0, 300 mM NaCl, and 10% glycerol supplemented with 2 mM of desthiobiotin. Elution fractions were analyzed by SDS-PAGE, and fractions containing the desired protein combined and concentrated using spin concentrators (Millipore). Following concentration, proteins were additionally purified via size exclusion chromatography (Cytiva HiLoad 26/600 Superdex-200 pg column, #GE28-9898-36). Pure proteins were concentrated, flash frozen in liquid nitrogen, and stored at −80 °C until further use.

ZF5.3-2D9 expression and purification

The plasmid encoding Strep-ZF5.3-2D9 was used to transform *E. coli* BL21 cells. Individual colonies were selected based on ampicillin resistance and used to inoculate 150 mL of LB media supplemented with ampicillin (100 µg/mL). The primary culture was grown overnight and then used to inoculate 6 L of LB media supplemented with ampicillin, which was then allowed to grow at 37 °C with shaking at 200 rpm. At $OD_{600nm}$ of 0.5, protein expression was induced by the addition of IPTG to a final concentration of 0.5 mM. After culturing for overnight at 18 °C, cells were harvested by centrifugation (6000 × g, 30 min at 4 °C), resuspended in buffer containing 25 mM Tris-HCl pH 8.0, 300 mM NaCl, 0.1 mM $ZnSO_4$, and 10% glycerol, supplemented with 1 mM PMSF, and lysed by microfluidization. The purification of this protein was identical to that of 2D9, with the exception that 0.1 mM of $ZnSO_4$ was added to all buffers.

ZF5.3-2D9-SPOP expression and purification

The plasmid encoding $His_6$-ZF5.3-2D9-SPOP was used to transform *E. coli* BL21 cells. Individual colonies were selected based on kanamycin resistance and used to inoculate 150 mL of LB media supplemented with kanamycin (50 µg/mL). The primary culture was grown overnight and then used to inoculate 6 L of LB media supplemented with kanamycin, which was then allowed to grow at 37 °C with shaking at 200 rpm. At $OD_{600nm}$ of 0.5, protein expression was induced by the addition of IPTG to a final concentration of 0.5 mM. After culturing for overnight at 18 °C, cells were harvested by centrifugation at 6000 × g for 30 min at 4 °C), resuspended in buffer containing 25 mM Tris-HCl pH 8.0, 300 mM NaCl, and 10% glycerol, supplemented with 1 mM PMSF, and lysed by microfluidization. The lysate was centrifuged at 15,000 × g for 45 min at 4 °C, and to the clear lysate was added 8 M urea, pH 8.0. The lysate was manually added to a column with 3 mL Ni-NTA resin and washed with 300 mL of buffer containing 25 mM Tris-HCl pH 8.0, 300 mM NaCl, 20 mM imidazole, and 8M urea. After that, protein was eluted from the resin using 25 mM Tris-HCl pH 8.0, 300 mM NaCl, 8M urea supplemented with 100 mM of imidazole. The elution fractions were analyzed by SDS page and concentrated to 1 mL. The purified proteins were refolded with 100 mL buffer containing 25mM Tris-HCl pH 8.0, 300 mM NaCl, 500 mM L-arginine, 9 mM glutathione and 1 mM glutathione disulfide. The refolded protein was cleared by centrifugation at 15,000 × g for 45 min at 4 °C, passed through a 0.22 μm filter, and concentrated for future use.

ZF5.3-2D9-RNF4 expression and purification

The plasmid encoding Strep-ZF5.3-2D9-RNF4 was used to transform *E. coli* BL21 cells. Individual colonies were selected based on ampicillin resistance and used to inoculate 150 mL of LB media supplemented with kanamycin (100 μg/mL). The primary culture was grown overnight and then used to inoculate 6 L of LB media supplemented with kanamycin, which was then allowed to grow at 37 °C with shaking at 200 rpm. At $OD_{600nm}$ of 0.5, protein expression was induced by the addition of IPTG to a final concentration of 0.5 mM. After culturing overnight, cells were harvested via centrifugation, resuspended in buffer containing 25 mM Tris-HCl pH 8.0, 300 mM NaCl, and 10% glycerol, supplemented with 1 mM PMSF, and lysed by microfluidization. The inclusion bodies were obtained by centrifugation at 15,000 × g for 45 min at 4 °C, and solubilized in 300 mM NaCl, 25 mM Tris-HCl pH 8.0, and 8 M urea. After another round of centrifugation at 15,000 × g for 45 min at 4 °C, the solubilized inclusion bodies (~2 mL) were mixed dropwise with 200 mL of refolding buffer containing 300 mM NaCl, 25 mM Tris-HCl pH 8.0, 500 mM L-arginine, 9 mM glutathione and 1 mM glutathione disulfide. The refolded mixture was cleared by centrifugation at 15,000 × g for 45 min at 4 °C and passed through a 0.22 μm filter. The refolded proteins were concentrated and saved for future use.

ZF5.3-SPOP expression and purification

The plasmid encoding $His_6$-ZF5.3-SPOP was used to transform *E. coli* BL21 cells. The expression and purification of this protein was identical to that of ZF5.3-2D9-SPOP.

***Protein binding assays***. Purified $His_6$-ZnF23, $His_6$-exZnF23 of BCL11A and $His_6$-exZnF23 of BCL11B were immobilized on the $Ni^{2+}$-NTA resin (Qiagen) equilibrated with the binding buffer (25mM Tris-HCl pH 8.0, 150mM NaCl, 1mM DTT). After 1h incubation at 4°C, beads were washed with the binding buffer. Purified nanobody proteins were then added to the beads and incubated at 4°C for another 1h. Beads were washed for at least three times with the binding buffer. The bound proteins were separated with SDS-PAGE and stained with Coomassie blue.

***Molecular mass analysis***. Molecular masses were analyzed by SEC-MALS with miniDawn Multi-Angle Light Scattering (MALS) detector, Optilab refractive index detector (Wyatt Technology, Santa Barbara, CA, USA) and UV (Waters 2487, Waters Corporation, Milford, MA) detectors. Volumes of injection is 15 µl (about 200 ng for each sample). The proteins were centrifuged at 13000 rpm for 10 min and then applied to pre-equilibrated Superdex 200 Increase 3.2/300 GL SEC column (Cytiva) with buffer containing 25 mM Tris-HCl, pH 8.0, 300 mM NaCl, 100 µM $ZnSO_4$ and 7 mM BME. Proteins were separated using a mobile phase flow rate of 0.15 mL/min at room temperature. Molecular masses were calculated using the Astra software (version 6) provided with the instrument.

***Microscale thermophoresis assay***. In vitro purified $His_6$-exZnF23 of BCL11A at a concentration of 1mg/mL was labeled with Alexa Fluor 647 dye according to the manufacturer's instruction (Invitrogen). After labeling, 100 nM BCL11A exZnF23 in buffer containing 25mM HEPES and 150 mM NaCl was used. 25 µL of 2D9 or ZF5.3-2D9 in the same buffer at the concentration of 67 and 22.6 µM, respectively, were used for 1:1 serial dilution into PCR-tubes. For each sample, 10 µL of labelled BCL11A exZnF23 was mixed with 10 µL ligand. Samples were loaded into capillaries and measured by MST with 40% LED power and medium MST power. Data was analyzed by the software MO Affinity Analysis (NanoTemper) using the "$K_d$" model and plotted in Prism. The equation below was used for fitting:

$$f(Concentration) = Unbound + \frac{(Bound - Unbound) * (Concentration + TargetConc + Kd - \sqrt{(Concentration + TargetConc + Kd)^2 - 4Concentration * TargetCon}}{2 * TargetConc})$$

Unbound: Response value of unbound state
Bound: Response value of bound state
TargetConc: final concentration of fluorescent molecules

***Crystallization of nanobody***. Sparse-matrix crystallization screens were performed with purified proteins at concentrations of about 11 mg/ml. Crystals were obtained by the sitting-drop vapor diffusion method by mixing 0.5 μL protein and 0.5 μL precipitant solution at 20°C. 2D9 nanobody crystals appeared in the SaltRX-F3 condition containing 1.5 M Ammonium sulfate, 0.1 M Tris-HCl, and pH 8.5 after 4 weeks of incubation at 20 °C. Crystals were cryoprotected with paraffin oil before being flash-cooled and stored in liquid nitrogen.

***Crystal data collection, processing, and structure determination***. All diffraction datasets were collected at 100 K at the Stanford Synchrotron Radiation Lightsource (Menlo Park, CA, USA). 2D9 datasets were collected at a fixed wavelength at 0.97946 Å, using an Eiger 16M detector at beamline BL12-1. All diffraction datasets were indexed and processed by XDS (*37, 38*). The structure was phased by molecular replacement in Phenix (Phaser) using model of 1NLB-H (*39*). Subsequent density modification gave excellent electron-density maps, which allowed the building of the model. Iterations of refinement were carried out with Phenix Refine, and model building was performed in Coot. The statistics for data collection, processing, and refinement are listed in Table S3. All structural figures were prepared using PyMOL.

***Western blot.*** After treatment, cells were washed with PBS buffer and lysed using lysis buffer (1% SDS, 150 mM NaCl, 0.1 U Benzonase nuclease (Santa Cruz biotechnology, #sc-391121B), EDTA-free protease inhibitor cocktail (Bimake, #B14002), 20 mM Tris-HCl, pH 8.0). Protein extracts were quantified by Pierce BCA Protein Assay Kit (Thermo Scientific, #PI23235) with a Nanodrop One$^{C}$ (Thermo Scientific) according to the manufacturer's instruction. Protein lysates (~ 50 μg/lane) were resolved by SDS-PAGE and transferred onto poly (vinylidene difluoride) (PVDF) membranes. Membranes were incubated with 5% non-fat milk in 0.1% Tween 20/PBS for 1 h. The blots were then probed with the relevant primary antibodies in blocking solution at 4°C overnight with gentle agitation. Membranes were washed 5 min with 0.1% Tween 20 in PBS three times and were incubated with Horseradish Peroxidase (HRP)-conjugated secondary antibody for 1 h at room temperature. Antigens were detected by addition of Clarity™ Western ECL Substrate (Biorad, #1705060). The membranes were visualized by chemiluminescence on a GE Amersham Imager

600. Antibodies used: Anti-BCL11A (Santa Cruz biotechnologies, #sc-514842), StrepMAB-Classic HRP (IBA, #2-1509-001), Anti-His-HRP (Sigma-Aldrich, #A7058), Anti-GAPDH (Santa Cruz Biotechnology, #sc-365062), Anti-Lamin B1 (Proteintech, #66095-1), Anti-β hemoglobin (Sigma Aldrich, #WH0003043M1), Anti γ-hemoglobin (Proteintech, #25728-1-AP). Secondary antibodies used: Anti-Mouse-HRP (Abcam, #ab6728), Anti-Rabbit-HRP (Promega, #W4018).

***BCL11A co-immunoprecipitation from HUDEP-2 cell lysate***. $8\times10^6$ HUDEP-2 cells was lysed with 400 µL cell lysis buffer (containing 25 mM Tris, 150 mM NaCl, 1% Triton-X100, 1% protease inhibitor cocktail, and 0.1 % SDS) for 10 min on ice to obtain the cell lysate. To 180 µg of total protein in the cell lysate was added 20 µg pure Nb proteins or buffer (as control). The cell lysate was first incubated for 3 h at 4 °C with gentle agitation and then incubated with 100 µL Strep-Tactin® Sepharose® for 4h at 4 °C. After that, resin was centrifuged (1000 × g, 2 min), and supernatant was carefully collected as flow-through. The resin was then washed with lysis buffer, centrifuged (1000 × g for 2 min), and the supernatant removed carefully. Finally, resin was incubated with elution buffer containing 2.5 mM desthiobiotin, centrifuged, and the eluate collected in the supernatant. Protein input and the eluate were immunoblotted with StrepMAB-Classic HRP and anti-BCL11A antibodies.

***Protein delivery.*** $4 \times 10^5$ cells were washed with PBS twice and seeded into 24-well plate in 500 µL serum free medium or medium containing proteins (as indicated in figures) for 45 min at 37 °C. After this, the cells were collected via by centrifugation (300 × g for 5 min) and incubated in full expansion medium at 37 °C for 45 min (or as indicated in the figures). Finally, the cells were harvested by centrifuge at 300 × g for 5 min and lysed for western blot.

***Nucleus isolation***. To isolate the nuclei, HUDEP-2 cells were first lysed with lysis buffer containing 25 mM Tris, 150 mM NaCl, 1% Triton-X100, 1% protease inhibitor cocktail, and 0.1% SDS for 10 min on ice. After this, cells were spun down at the maximum speed to get the nuclei at the bottom as white pellets. The supernatant was collected as cytosolic fractions. The nucleus fraction was then washed three times with PBS buffer and lysed with buffer containing 25 mM Tris, 1% SDS, 150 mM NaCl, 0.1% benzonase, and 2 mM EDTA to obtain the nuclear proteins.

***Confocal imaging***. ZF5.3-2D9V102G at the concentration of 10 μM was delivered into $4 \times 10^5$ HUDEP-2 cells, and cells were cultured for another 24 h in full expansion medium at 37 $^{o}$C. After that, cells were washed with PBS for 3 times, and seeded into confocal plates coated with poly-lysine (Fisher Scientific, #80824). Cells were cultured at 37 $^{o}$C for 1h to allow settling down after seeding. Cells were then and fixed by addition of 4% paraformaldehyde for 20 min at 4 $^{o}$C and permeabilized by adding 0.1% Triton-X100 in PBS for 5 min. After staining with Anti-Strep, and Anti-Rab7 (Cell Signaling, #9367) at 4 $^{o}$C overnight, fixed cells were washed with 0.1% Tween 20 in PBS for 3 times, cells were then incubated with 1 μg/mL DAPI, secondary antibodies Goat anti-rabbit IgG (H+L) -488 (Thermo Scientific, #A32371) and Goat anti-Mouse-IgG (H+L) -Alexa Fluor 647 (Thermo Scientific, #A21235) for 1h at room temperature. Cells were washed 3 times with Tween 20 in PBS and imaged with a Zeiss LSM980 with Airyscan 2 Confocal Microscope with 60 × oil immersion objective at excitations of 653, 493 and 353 nm.

***AlphaScreen assay.*** The assay was performed in a light grey 384-well AlphaPlate (PerkinElmer) containing 10 nM biotinylated Avi-tagged nanobody protein and a serial dilution of $His_6$-tagged exZnF23 of BCL11A in a total volume of 20 μL in AlphaScreen buffer (20 mM Tris-HCl pH 7.5, 150 mM NaCl, mM DTT, 0.1% Tween, 0.1% BSA). Reaction mixtures were incubated at RT for 30 min. Streptavidin Donor beads and nickel chelate (Ni-NTA) AlphaScreen Acceptor beads (PerkinElmer) each at a concentration of 10 μg/mL were added to the mixture. The plate was incubated at RT for 1 hr in the dark and then analyzed by a plate reader.

***Transduction or transfection of Nb-Fc, Nb-Trim21 and Nb-mutTrim21***. To generate nanobody-Fc, nanobody-wtTrim21 and nanobody-mutTrim21 fusions, the nanobody coding sequence followed by the hIgG1-Fc coding sequence (pFuse-hIgG1-Fc1; Invivogen), or full-length wild-type Trim21 or mutant Trim21 (M10E/M72E) were subcloned into the lentiviral vector pLVX-EF1a-IRES-ZsGreen (Clontech #631982). Lentiviruses were packaged in HEK293T cells as described previously (*40*).

For expression in HEK293T cells, plasmids were transfected using Lipofectamine 2000 (Invitrogen) according to manufacturer's instructions. Cells were harvested 24 hours after transfection, subjected to SDS-PAGE and analyzed by Western Blotting.

HUDEP-2 cells were transduced with lentiviruses at a confluency of $2\times10^5$ cells/mL. GFP positive cells were FACS sorted 72hr post-transduction. After sorting, cells were collected at Day 7 after differentiation for Western Blotting and RT-qPCR.

***RT-qPCR***. Quantitative real-time PCR Quantitative real-time PCR was performed to quantify RNA abundance. For each sample, total RNA was isolated by using the RNeasy Plus Mini Kit (QIAGEN Cat# 74134), followed by cDNA synthesis using the iScript cDNA Synthesis Kit (BioRad, Cat# 1708890). qRT-PCR primers (Table S4) were ordered from Integrated DNA Technologies. Quantitative real-time PCR was performed using the PrimePCR assay with the iTaq universal SYBR Green supermix (BioRad, #1725120) and run on a Biorad CFX384 real-time system (C1000 Touch Thermal Cycler) according to the manufacturer instructions. $C_q$ values (Tables S5, S6) were used to quantify RNA abundance. The relative abundance of the hemoglobin was normalized to a GAPDH internal control by using this equation:

$$\Delta Cq = Cq\ (\text{gene of interest}) - Cq\ (\text{GAPDH})$$

$$R = 2^{-\Delta Cq}$$

***Fluorescence Activated Cell Sorting (FACS)***. Cells were harvested and washed twice with PBS buffer and centrifuged at 350-500 × g for 5 min. Then, cells were fixed with 4% paraformaldehyde for 20 min at 4 °C and permeabilized with 0.2 % Triton-X100 in PBS for 5 min. Following that, 100 µL of 0.1% BSA in 0.1% Tween 20/PBS buffer was added for blocking, and cells were incubated with anti γ-hemoglobin (Proteintech, #25728-1-AP) for 2 h, washed 3 times with 0.1% Tween 20 in PBS buffer and stained with Goat anti-rabbit IgG (H+L) -488 (Thermo Scientific, #A32371) for 1 h at room temperature. After washing 3 times with 0.1% Tween 20 in PBS buffer, cells were analyzed on a BD Accuri™ C6 Plus cell sorter.

### Supplemental figures

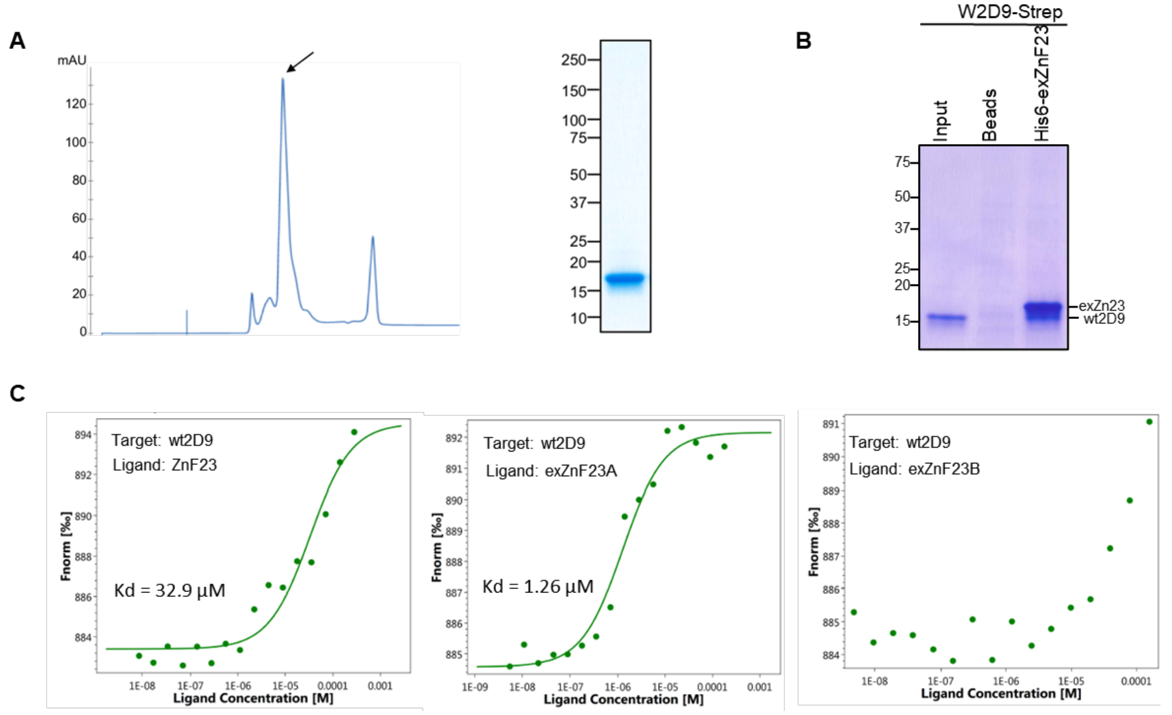


**Figure S1**. (**A**) Size-exclusion chromatogram and SDS-PAGE analysis of purified wt2D9 (arrow pointing to fraction with pure protein). (**B**) SDS-PAGE gel showing the pull-down of BCL11A exZnF23 by wt2D9. (**C**) MST binding curves of wt2D9 to BCL11A ZnF23 (left), BCL11A exZnF23 (exZnF23A, middle) and BCL11B exZnF23 (exZnF23B, right).

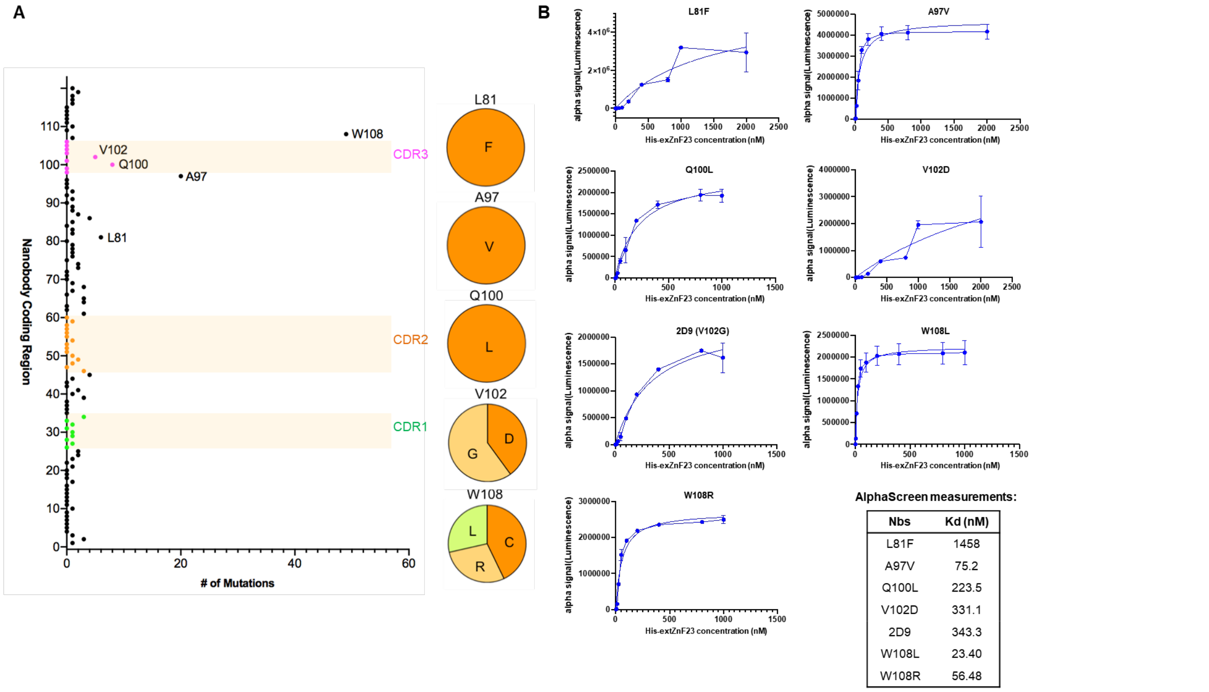


| Nbs | Kd (nM) |
|---|---|
| L81F | 1458 |
| A97V | 75.2 |
| Q100L | 223.5 |
| V102D | 331.1 |
| 2D9 | 343.3 |
| W108L | 23.40 |
| W108R | 56.48 |

**Figure S2.** (**A**) Identification of maturated nanobodies targeting exZnF23 of BCL11A. Sequence analysis of 96 randomly picked yeast colonies following MACS and FACS enrichment reveals mutations that favor binding to exZnF23 of BCL11A. (**B**) The binding affinities of maturated nanobodies to exZnF23 of BCL11A were measured by AlphaScreen.

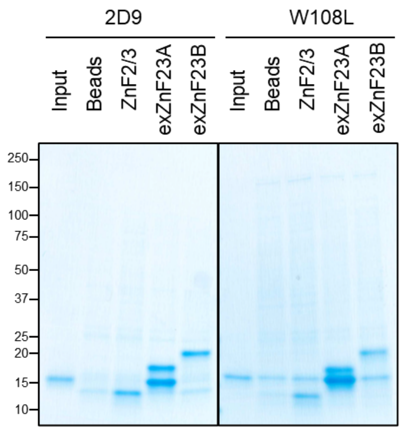


**Figure S3**. SDS gels showing that 2D9 and W108L pull down BCL11A exZnF23 (exZnF23A) but not BCL11B exZnF23 (exZnF23B).

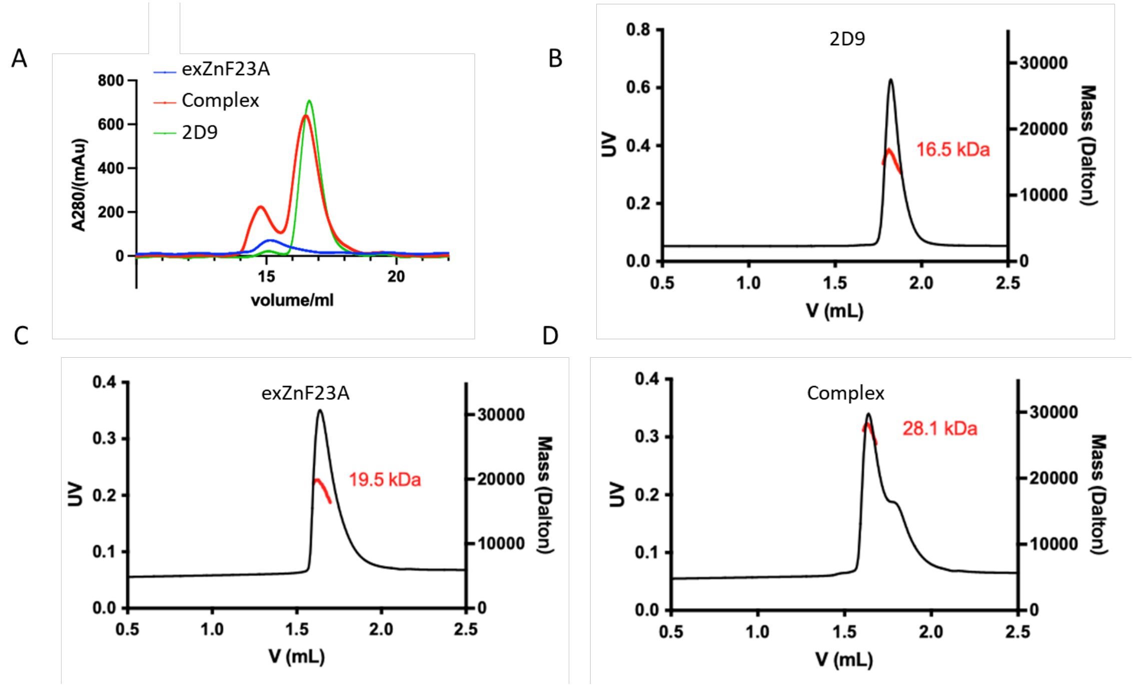


**Figure S4**. **(A)** Analytical size-exclusion chromatography (SEC) of BCL11A exZnF23, 2D9, and the complex they form. 100 nM of purified exZnF23, 2D9, and the mixture of the two at a 1:1 molar ratio (pre-incubated for 0.5 h on ice) was injected into Superdex 200 10/300GL and eluted in buffer containing 25 mM Tris-HCl pH 8.0, 300 mM NaCl, 100 µM $ZnSO_4$ and 7 mM β-mercaptoethanol. SEC-MALS analysis of **(B)** 2D9, **(C)** exZnF23, and **(D)** their complex revealed that exZnF23 is monomeric and forms a stable complex with 2D9.

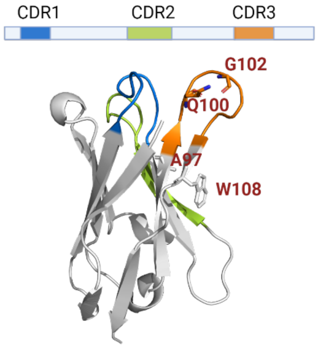


**Figure S5**. Cartoon depiction of the crystal structure of 2D9. CDR3 region is highlighted in green. Mutations of the Nb (A97, Q100, G102and W108) that produced proteins with improved binding to BCL11A are highlighted in red.

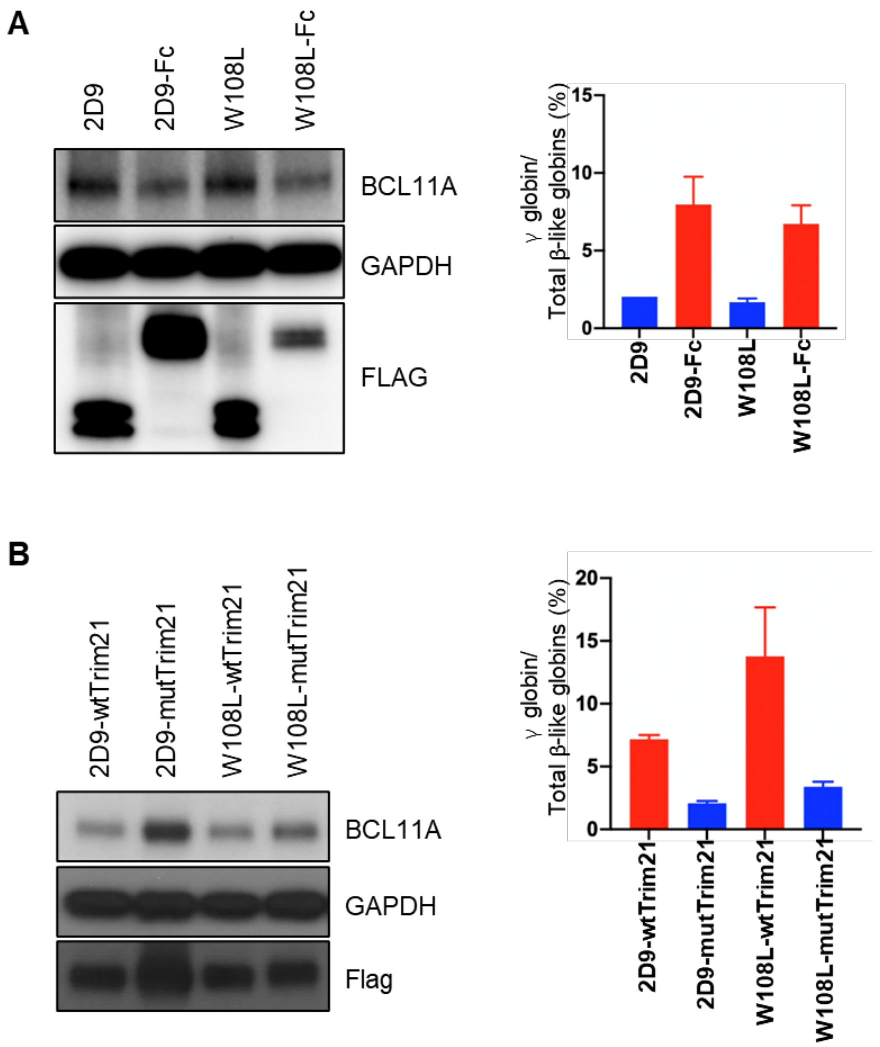


**Figure S6.** (**A**) Degradation of BCL11A and reactivation of γ-globin upon lentiviral delivery of Nb-Fc fusion in differentiated HUDEP-2 cells. (**B**) HUDEP-2 cells were transduced with the indicated nanobodies fused with wild-type Trim21 or catalytically inactive (mut)Trim21. Lentiviral delivery of Nb-wtTrim21 but not Nb-mutTrim21 led to degradation of BCL11A and reactivation of γ-globin in differentiated cells.

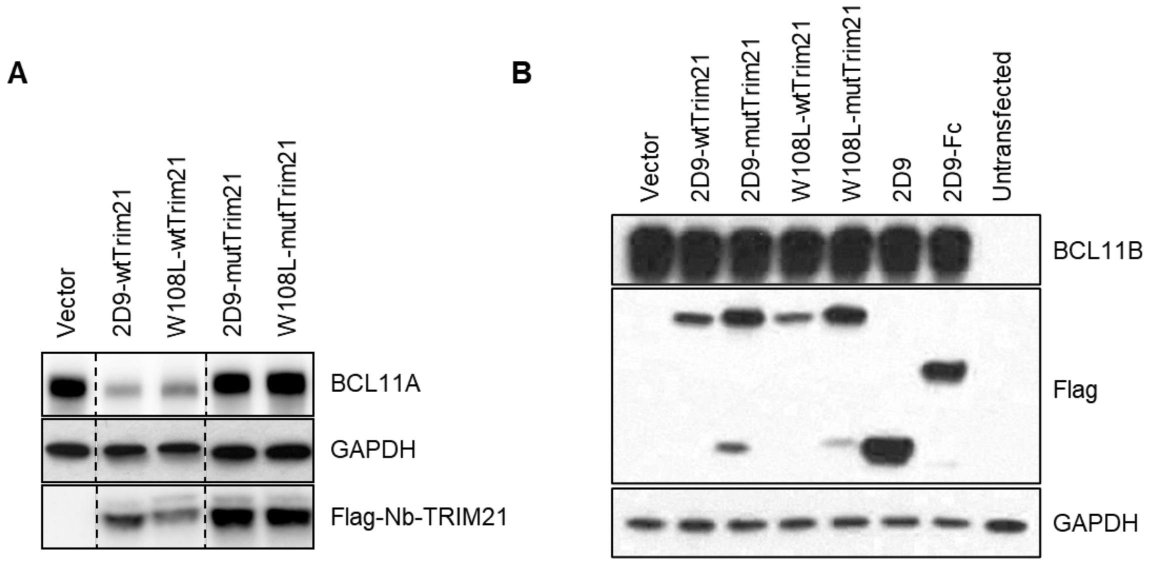


**Figure S7**. (**A**) Nb-wtTrim21 but not Nb-mutTrim21 led to degradation of overexpressed BCL11A in HEK293T cells. (**B**) Immunoblotting shows that neither Nb-Fc nor Nb-Trim 21 mediated degradation of overexpressed BCL11B in HEK293T cells.

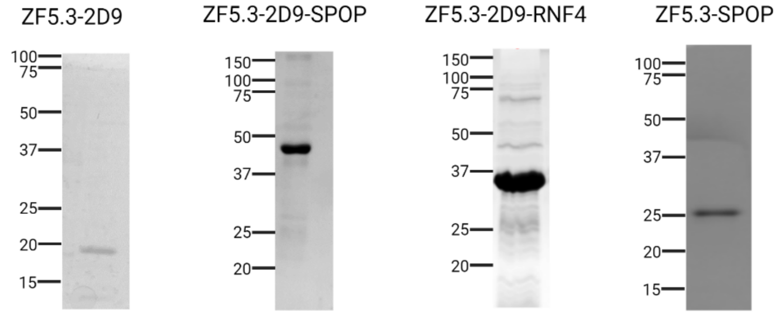


**Figure S8**. SDS-PAGE gels of purified proteins.

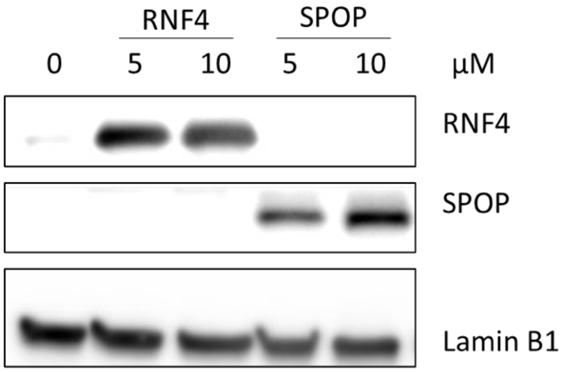


**Figure S9**. Immunoblotting of HUDEP-2 cell lysate reveals the cellular uptake of both of ZF5.3-2D9-SPOP and ZF5.3-2D9-RNF4.

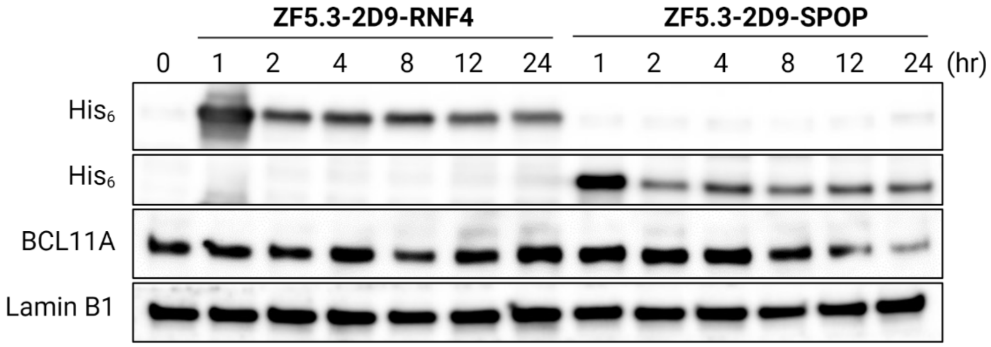


Figure **S10**. Immunoblot showing loss of BCL11A in HUDEP-2 cells treated with ZF5.3-2D9-RNF4 or ZF5.3-2D9-SPOP. This is a second biological replicate of the data shown in Figure 3.

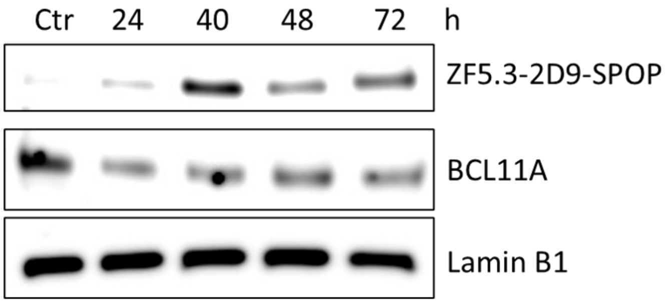


**Figure S11**. Persistence of BCL11A degradation in HUDEP-2 cells. HUDEP-2 cells were incubated with 10 µM ZF5.3-2D9-SPOP for 45 min, washed, and cultured in fresh medium without ZF5.3-2D9-SPOP for the times as indicated.

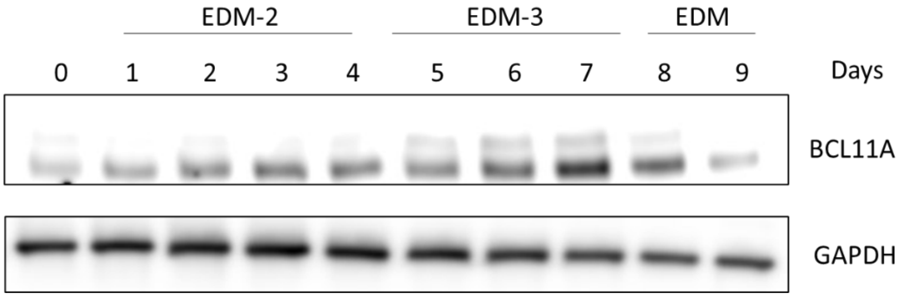


**Figure S12**. BCL11A levels in differentiated HUDEP-2 cells. Upon differentiation, BCL11A levels gradually increase and peak at Day 7 before decreasing to basal levels.

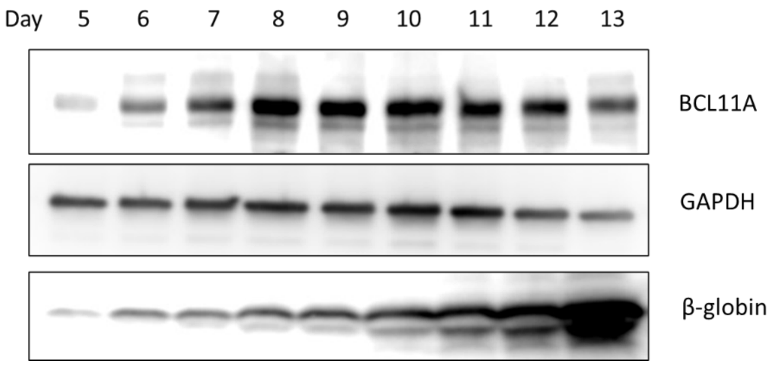


**Figure S13**. BCL11A and β-hemoglobin levels in differentiated $CD34^+$ cells. Upon differentiation, BCL11A levels gradually increased, peaked at Day 8, and then decreased to basal levels.

| Table S1. Primers used for plasmid construction | | |
|---|---|---|
| **Plasmid** | **Primer** | **Sequence (5’-3’)** |
| pET-20b_ZF5.3-2D9 | pET-20b 2D9 linearization-FWD | AGCGGGCAAGTTCAGCTGGTTG |
| | pET-20b 2D9 linearization-FWD | GCTGTAGGATCCCATGTGACC |
| | ZF5.3-Insertion-FWD | CAGGGTCACATGGGATCCTACAGCTGCAACGTG |
| | ZF5.3-Insertion-REV | CTCAACCAGCTGAACTTGCCCGCTACCAGTCG |
| pET-20b_ZF5.3-2D9-SPOP | pET-20b_2D9 linearization-FWD | TAACTCGAGCACCAC |
| | pET-20b_2D9 linearization-FWD | TGAACCGCTGCCGCTGGAAACGGTAAC |
| | SPOP-insertion-FWD | GCGGCAGCGGTTCATCAGTAAATATCTCAGGGC |
| | SPOP-insertion-REV | GTGCTCGAGTTAGGACTGTTTGAGACTTTG |
| pET-20b_ZF5.3-2D9-RNF4 | RNF4-insertion-FWD | GCGGCAGCGGTTCAGAGGAACGGCGTC |
| | RNF4-insertion-REV | GTGCTCGAGTTAAATGTAAATCGGATGATAACG |
| pET-28a_ZF5.3-SPOP | pET-28a_ZF5.3-2D9-SPOP Linearization-FWD | TAACTCGAGCACCAC |
| | pET-28a_ZF5.3-2D9 -SPOP Linearization-REV | ATGAACCGCTGCCAGTCGCGCGACGAT |
| | Insertion-FWD | ACTGGCAGCGGTTCATCAGTAAATATCTCAGGGC |
| | Insertion-REV | GGTGGTGCTCGAGTTAGGACTGTTTGAGACG |

| Table S2. Complete sequence of proteins used for the cell-permeant degrader | |
|---|---|
| **Protein** | **Amino acid sequence** |
| wt2D9 | QVQLVESGGGLVQAGGSLRLSCAASGSIFVNNAMGWYRQAPGKERELVAAISASGGSTYYADSVKGRF TISRDNAKNTVYLQMNSLKPEDTAVYYCAADQDVYPYEYWGQGTQVTVS |
| ZF5.3-2D9 | MLESAWSHPQFEKENLYFQGHMGSYSCNVCGKAFVLSRHLNRHLRVHRRATGSGQVQLVESGGGLVQ AGGSLRLSCAASGSIFVNNAMGWYRQAPGKERELVAAISASGGSTYYADSVKGRFTISRDNAKNTVYLQ MNSLKPEDTAVYYCAADQDVYPYEYWGQGTQVTVSS |
| ZF5.3-2D9-SPOP | MGSSHHHHHHSSGLVPRGSHMYSCNVCGKAFVLSRHLNRHLRVHRRATGSGQVQLVESGGGLVQAG GSLRLSCAASGSIFVNNAMGWYRQAPGKERELVAAISASGGSTYYADSVKGRFTISRDNAKNTVYLQMN SLKPEDTAVYYCAADQDGYPYEYWGQGTQVTVSSGSGSSVNISGQNTMNMVKVPECRLADELGGLWE NSRFTDCCLCVAGQEFQAHKAILAARSPVFSAMFEHEMEESKKNRVEINDVEPEVFKEMMCFIYTGKAP NLDKMADDLLAAADKYALERLKVMCEDALCSNLSVENAAEILILADLHSADQLKTQAVDFINYHASDVLE TSGWKSMVVSHPHLVAEAYRSLASAQCPFLGPPRKRLKQS |
| ZF5.3-SPOP | MGSSHHHHHHSSGLVPRGSHMYSCNVCGKAFVLSRHLNRHLRVHRRATGSGSSVNISGQNTMNMVK VPECRLADELGGLWENSRFTDCCLCVAGQEFQAHKAILAARSPVFSAMFEHEMEESKKNRVEINDVEPE VFKEMMCFIYTGKAPNLDKMADDLLAAADKYALERLKVMCEDALCSNLSVENAAEILILADLHSADQLKT QAVDFINYHASDVLETSGWKSMVVSHPHLVAEAYRSLASAQCPFLGPPRKRLKQS |
| ZF5.3-2D9-RNF4 | MLESAWSHPQFEKENLYFQGHMGSYSCNVCGKAFVLSRHLNRHLRVHRRATGSGQVQLVESGGGLVQ AGGSLRLSCAASGSIFVNNAMGWYRQAPGKERELVAAISASGGSTYYADSVKGRFTISRDNAKNTVYLQ MNSLKPEDTAVYYCAADQDGYPYEYWGQGTQVTVSSGSGSEERRRPRRNGRRLRQDHADSCVVSSDD EELSKDKDVYVTTHTPRSTKDEGTTGLRPSGTVSCPICMDGYSEIVQNGRLIVSTECGHVFCSQCLRDSLK NANTCPTCRKKINHKRYHPIYI |

| Table S3. Crystallographic data collection and refinement statistics for 2D9 | |
|---|---|
| Resolution range (Å) | 30.44 - 1.25 (1.295 - 1.25) |
| Space group | P 21 21 21 |
| Unit cell: a, b, c (Å); $\alpha$, $\beta$, $\gamma$ (°) | 36.619, 52.247, 54.757<br>90°, 90°, 90° |
| Total reflections | 213480 (9759) |
| Unique reflections | 29034 (2441) |
| Multiplicity | 7.4 (4.0) |
| Completeness (%) | 97.59 (83.50) |
| Mean I/sigma | 17.70 (1.82) |
| Wilson B-factor | 15.57 |

| | |
|---|---|
| R-merge | 0.0703 (0.4882) |
| R-meas | 0.07538 (0.5652) |
| R-pim | 0.02671 (0.2716) |
| CC1/2 | 0.999 (0.834) |
| CC* | 1 (0.954) |
| Reflections used in refinement | 29027 (2440) |
| Reflections used for R-free | 1991 (162) |
| R-work | 0.1974 (0.3003) |
| R-free | 0.2159 (0.3115) |
| CC(work) | 0.962 (0.870) |
| CC(free) | 0.940 (0.864) |
| Number of non-hydrogen atoms | 1026 |
| macromolecules | 935 |
| ligands | 10 |
| solvent | 81 |
| Protein residues | 124 |
| RMS(bonds) | 0.012 |
| RMS(angles) | 1.55 |
| Ramachandran favored (%) | 97.54 |
| Ramachandran allowed (%) | 2.46 |
| Ramachandran outliers (%) | 0.00 |
| Rotamer outliers (%) | 1.04 |

| | |
|---|---|
| Clashscore | 2.19 |
| Average B-factor | 19.83 |
| macromolecules | 19.03 |
| ligands | 38.47 |
| solvent | 26.68 |

| **Table S4. Primes used for RT-qPCR** | |
|---|---|
| HBB-FWD: | CTGAGGAGAAGTCTGCCGTTA |
| HBB- REV: | AGCATCAGGAGTGGACAGAT |
| HBE- FWD: | GCAAGAAGGTGCTGACTTCC |
| HBE- REV | ACCATCACGTTACCCAGGAG |
| HBG- FWD: | TGGATGATCTCAAGGGCAC |
| HBG- REV: | TCAGTGGTATCTGGAGGACA |
| GAPDH-FWD: | ACCCAGAAGACTGTGGATGG |
| GAPDH-REV: | TTCAGCTCAGGGATGACCTT |

**Table S5. $C_q$ values of hemoglobin transcripts in HUDEP-2 cells treated with ZF5.3-2D9-SPOP**

| HUDEP-2 cells (Day 7) | | | | | | | | | | | | |
|---|---|---|---|---|---|---|---|---|---|---|---|---|
| | Untreated | | | | | | Treated | | | | | |
| | Cq1 | Cq 2 | Cq 3 | Cq Avg | Δ Cq | $2^{-\Delta Cq}$ | Cq 1 | Cq 2 | Cq3 | Cq Avg | Δ Cq | $2^{-\Delta Cq}$ |
| HBB | 12.33 | 12.28 | 12.65 | 12.42 | -10.31 | 1272.39 | 14.43 | 14.34 | 14.77 | 14.51 | -10.84 | 1833.01 |
| HBG | 21.68 | 21.59 | 22.15 | 21.81 | -0.93 | 1.90 | 22.20 | 22.04 | 22.91 | 22.38 | -2.97 | 7.84 |
| GAPDH | 22.58 | 22.60 | 23.02 | 22.73 | / | / | 25.26 | 25.39 | 25.41 | 25.35 | / | / |

| Table S6. $C_q$ values of hemoglobin transcripts in $CD34^+$ cells treated with ZF5.3-2D9-SPOP | | | | | | | | | | | | |
|---|---|---|---|---|---|---|---|---|---|---|---|---|
| $CD34^+$ cells (Day 13) | | | | | | | | | | | | |
| | Untreated | | | | | | Treated | | | | | |
| | Cq1 | Cq 2 | Cq 3 | Cq Avg | Δ Cq | $2^{-\Delta Cq}$ | Cq 1 | Cq 2 | Cq3 | Cq Avg | Δ Cq | $2^{-\Delta Cq}$ |
| HBB | 14.63 | 13.35 | 13.47 | 13.81 | -10.98 | 2014.60 | 16.02 | 15.35 | 15.44 | 15.60 | -11.45 | 2799.59 |
| HBG | 18.53 | 17.04 | 16.94 | 17.50 | -7.23 | 156.23 | 17.29 | 16.49 | 16.48 | 16.75 | -10.30 | 1258.60 |
| GAPDH | 25.14 | 24.62 | 24.63 | 24.80 | / | / | 27.67 | 26.57 | 26.92 | 27.05 | / | / |
